# The effects of *CDC45* mutations on DNA replication and genome stability

**DOI:** 10.1101/2024.10.28.620574

**Authors:** Milena Denkiewicz-Kruk, Deepali Chaudhry, Alina Krasilia, Malgorzata Jedrychowska, Iwona J. Fijalkowska, Michal Dmowski

**Affiliations:** Institute of Biochemistry and Biophysics Polish Academy of Sciences Pawinskiego 5a, 02-106 Warsaw, Poland

**Keywords:** *CDC45*, CMG helicase, polymerase zeta, DRIM, mismatch repair, DNA replication fidelity

## Abstract

Cdc45 is a non-catalytic subunit of the CMG helicase complex and is recruited to the autonomously replicating sequence at the onset of DNA replication. Cdc45 protein is required for the initiation of the DNA replication process as well as for the nascent DNA strand synthesis. It interacts with Mcm2 and Psf1 elements of CMG helicase, as well as with Sld3, and initiation factor, and Pol2, the catalytic subunit of DNA polymerase epsilon (Pol ε). In this study, we analyzed the effects of amino acid substitutions in Cdc45 regions involved in the interaction of this protein with Mcm2-7 (Cdc45-1), Psf1 (Cdc45-26), and Sld3 (Cdc45-25, Cdc45-35). We found that mutations in *CDC45* result in defective DNA replication. At the restrictive temperature, mutant cells are unable to initiate DNA replication, while in permissive conditions, they demonstrate DNA synthesis delay. Moreover, we observed increased mutation rates, mainly dependent on DNA polymerase zeta (Pol ζ), as well as increased incidence of replication errors. These findings contribute to our understanding of Cdc45’s function in eukaryotic cells. Changes in the cell functioning observed in this study, related to the defect in Cdc45 function, may be helpful in understanding some diseases associated with *CDC45*.

## Introduction

Faithful replication of genetic information is crucial for maintaining genome stability. It requires the coordinated action of many catalytic as well as non-catalytic proteins during the initiation, elongation, and termination of DNA replication. In the first step of initiation, the six-subunit protein complex Orc1-6 (origin recognition complex) binds to the ARS (autonomously replicating sequence) [1]. Then, Cdc6 and Cdt1 proteins attach two Mcm2-7 heterohexamers complexes (inactive helicase cores), which together form the pre-Replicative Complex (pre-RC) [2]. In parallel, the Sld3-Sld7 complex binds the Cdc45 protein [3]. After phosphorylation of the Mcm2-7 heterohexamer by DDK kinase (Dbf4-dependent kinase), the Sld3-Sld7-Cdc45 complex is recruited to the pre-RC complex [4,5]. Finally, under the control of CDKs, at the transition of G1/S phases of the cell cycle, the pre-Loading Complex (pre-LC) composed of Dpb11, Sld2, GINS, and DNA polymerase epsilon (Pol ε) binds to the pre-RC, resulting in the formation of the pre-Initiation Complex (pre-IC). After forming the pre-IC, the Dpb11, Sld2, Sld3, and Sld7 proteins dissociate from the complex. As a result, the active CMG helicase consists of eleven proteins: the Cdc45 protein, the Mcm2-7 heterohexamer, and the GINS heterotetramer (Psf1-3, Sld5) [6] which, through interaction with DNA polymerase ε (Pol ε), forms an active fifteen-subunit CMGE complex, which unwinds the DNA double helix (CMG helicase) and replicates the leading strand (Pol ε) [7]. During the unwinding of double-stranded DNA by the CMG helicase, Replication Protein A (RPA) binds to and protects the single-stranded DNA (ssDNA) [8].

The results of recent studies indicate that the non-catalytic subunits of the helicase serve functions beyond their structural role and are crucial for the proper functioning of the entire replisome. Although the Cdc45 protein and the GINS complex are auxiliary subunits and have no catalytic activity, they play a key role in activating the CMG helicase [6,9]. In addition, Cdc45-GINS stabilizes the correct spatial structure of the Mcm2-7 complex and positions the leading DNA strand in the central channel, also preventing it from sliding out the Mcm2/5 gate in situations such as replication stress [10–12].

Early studies have shown that Cdc45 is essential for cell viability [13,14]. Later, it was shown that both yeast and human Cdc45 proteins bind to single-stranded DNA, which, under replication stress, activates the cellular response [15]. Initially, bioinformatic analyses have shown that Cdc45 is an orthologue of bacterial RecJ, a single-stranded DNA exonuclease belonging to the DHH family of phosphoesterases [16,17]. This was confirmed later when human and yeast Cdc45 structures were determined [18,19]. In general, Cdc45 contains DHH and DHHA1 domains separated by a connecting domain encompassing an unstructured region, a CID (CMG-interacting domain) for Mcm binding, and a RecJ fold involved in interaction with Psf1 [18,20] (Figure 1A).

**Figure 1.**
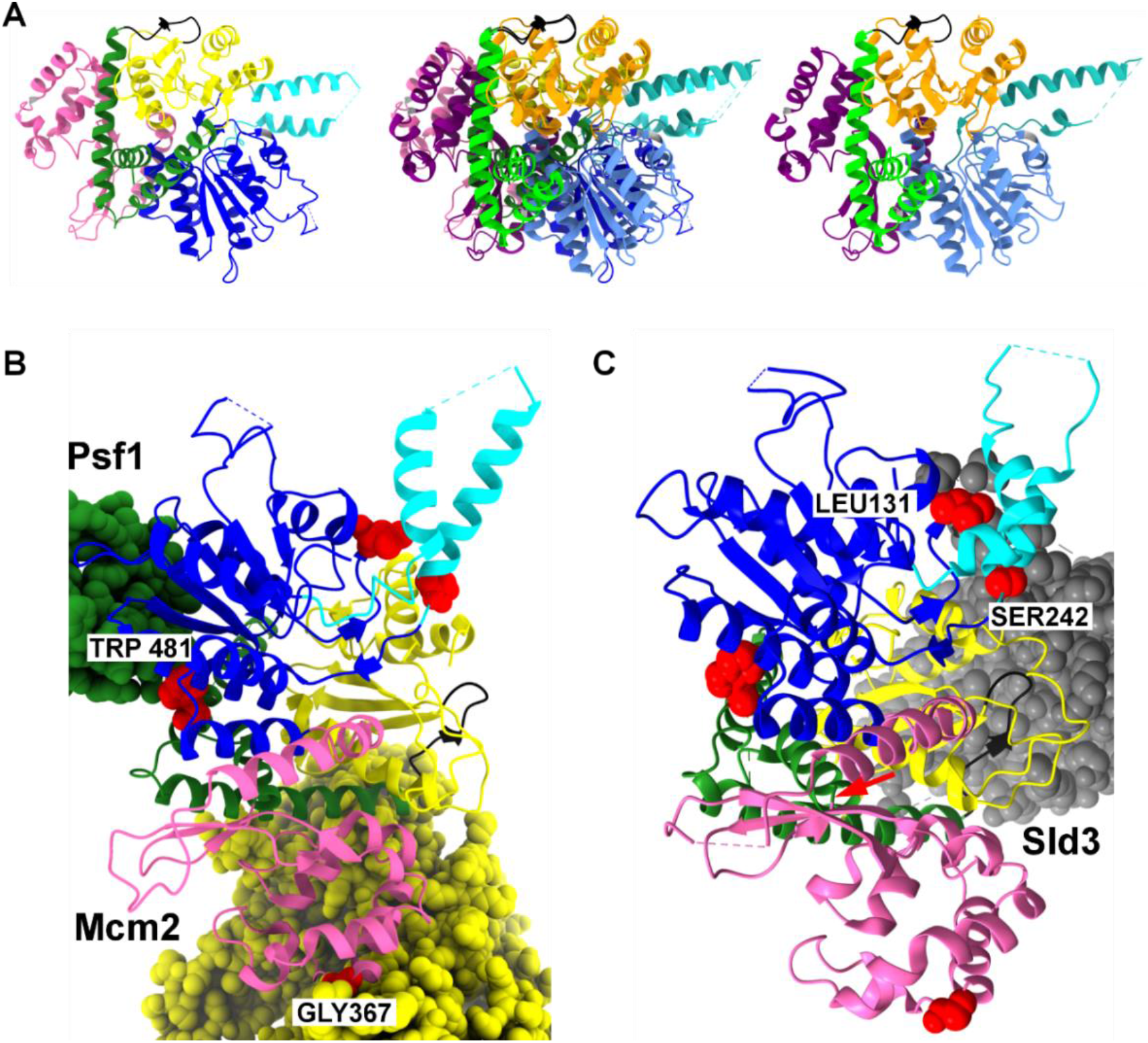
Structure of Cdc45: **A.** Yeast and human Cdc45 structures are shown separately (left and right) and overlayed (center) based on PDB 7qhs and 7pfo [62,63]. Protein domains are shown in specific colors: blue/cornflower blue – DHH; cyan/light sea green - helical insertion with unstructured region, absent in RecJ; green/lime – RecJ fold; pink/purple – CID; yellow/orange – DHHA1; **B.** Structure of Cdc45 and interacting proteins Mcm2 (yellow) and Psf1 (green) based on PDB 7qhs [62]; **C.** Structure of Cdc45 and Sld3 (grey) based on PDB Japan 8j09 [61]. Other components of CMG were omitted for clarity. Substituted residues in Cdc45 are shown as red spheres: G367D (Cdc45-1), L131P (Cdc45-25), W481R (Cdc45-26), and S242P (Cdc45-35). Structures according to PDB 7QHS [62] and PDB 7PFO [63].

Although numerous studies brought data on the structure of Cdc45 from various organisms (Figure 1A), a detailed analysis of the consequences of defective functioning of Cdc45 in eukaryotic cells is missing. In this study, we analyzed four alleles encoding Cdc45 variants with amino acid substitutions in the DHH domain L131P (Cdc45-25), S242P (Cdc45-35), CID domain (G367D) (Cdc45-1), and the Rec J fold W481R (Cdc45-26) (Fig 1B). One of them, *cdc45-1*, has been isolated previously [21].

We present data indicating that the proper functioning of Cdc45 in the replisome is important for genome stability. This conclusion is based on the analysis of four *cdc45* mutants that exhibit impaired progression through the S phase, elevated levels of replication errors, increased amounts of ssDNA, and more frequent contribution of the mutator Pol ζ to DNA synthesis.

Studies on the role of the Cdc45 protein are critical in the light of works describing that dysfunction of both alleles of the *CDC45* gene in humans leads to the development of an autosomal recessive disease – Meier-Gorlin syndrome (MGS) [22–24].

## Materials and Methods

### Strains, media, and general methods

The *Saccharomyces cerevisiae* strains were grown at 23°C or 30°C in standard media [25]. A complete YPD medium was used when nutrition selection was not required. YPD with nourseothricin 100 µg/mL (Werner BioAgents, Jena, Germany) and SD medium supplemented with appropriate amino acids and nitrogenous bases were used to select and propagate transformants. For the forward mutagenesis assays at the *CAN1* locus, the SD medium with appropriate amino acids and nitrogenous bases was supplemented with 60 µg/mL L-canavanine (Sigma Aldrich, St. Louis, MO, USA). The SD medium with 1 mg/mL 5-fluoroorotic acid (5-FOA) (US Biological, Salem, MA, USA) was used to select cells that lost the functional *URA3* gene.

Yeast cells were transformed using the lithium acetate/single-stranded carrier DNA/PEG method [26]. Chromosomal DNA was isolated from yeast cultures using the Genomic Mini AX Yeast Spin Kit (A&A Biotechnology, Gdansk, Poland).

*Escherichia coli DH5α* (*endA1 glnV44 thi-1 recA1 relA1 gyrA96 deoR nupG Φ80d lacZΔ M15Δ* (*lacZYA-argF*) U169, *hsdR17* (r K^−^,m K^+^), λ-) (Invitrogen, California, United States) cells were grown at 37 °C in LB medium, supplemented when needed with ampicillin 100 μg/mL (Polfa Tarchomin S.A., Warsaw, Poland). *E. coli* cells were transformed as described in [27]. Bacterial plasmids were isolated using the Plasmid Mini Kit (A&A Biotechnology, Gdansk, Poland).

### Construction of yeast strains

The *Saccharomyces cerevisiae* strains used in this study are derivatives of strain ΔI(- 2)I-7BYUNI300 [28], detailed in Table S1.

Yeast strains with mutated *CDC45* alleles are derivatives of SC765 obtained using integration cassettes derived from YIp211cdc45-1, YIp211cdc45-26 by BglII digestion, or YIp211cdc45-25, YIp211cdc45-35 by XbaI digestion. These plasmids, obtained from Hiroyuki Araki (NIG, Mishima, Japan), were constructed by subcloning 2.2 kb BamHI-HindIII fragments from YCp22CDC45 -derivatives with mutated alleles [29,30] into the BamHI-HindIII sites of YIplac211. The SC765 derivatives with *CDC45* alleles were selected on the SD medium without uracil. Transformants were grown at 30°C (*cdc45-1*) or 23°C (*cdc45-25, cdc45-26, cdc45-35*) for 5 to 7 days. Integrations of the *cdc45* alleles into the *CDC45* chromosomal locus were confirmed by PCR and DNA sequencing using primers cdc45_up, cdc45_dw, cdc45_1, cdc45_2, cdc45_3, cdc45_4 (Table S2). Additionally, temperature sensitivity at 37°C for *cdc45-25, cdc45- 26, and cdc45-35* strains and cold sensitivity at 18°C for the *cdc45-1* strain were verified. Finally, transformants were spread onto 5-FOA plates to select cells that had lost the *URA3* gene.

The *cdc45* strains (Y1050, Y1051, Y1052 and Y1053) and the *CDC45* control strain (SC765) were subjected to further modifications. For *MSH2* disruption, the *msh2::NAT1* disruption cassette was PCR-amplified with the primers MSH_UPTEF and MSH2_DNTEF (Table S2) using pAG25 [31] as a template. Transformants were grown on a YPD medium supplemented with nourseothricin (100 µg/mL) at 30°C (Y1050) or 23°C (Y1051, Y1052 and Y1053 strains) for 5 to 7 days. The integration of the of *msh2::NAT1* was confirmed by PCR using primers MSH2 A, MSH2 B, MSH2 C, MSH2 D, msh2_up, msh2_prdw, NAT1 UO and NAT1 DO (Table S2). For *REV3* disruption, the rev3::LEU2 disruption cassette, as described previously (Kraszewska et al. 2012), was used. Transformants were selected for leucine prototrophy on the SD medium at 30°C (Y1050) or 23°C (Y1051, Y1052, and Y1053 strains) for 5 to 7 days. The integration of *rev3::LEU2* was confirmed by PCR using primers REV3 A, REV3 B, REV3 C, and REV3 D (Table S2). The *RFA1-YFP* fusion was introduced as described previously [32]. Transformants were selected on the SD medium for leucine prototrophy at 30°C (Y1050) or 23°C (Y1051, Y1052, and Y1053 strains) for 5 to 7 days and verified by PCR, using the primers RFA6231R, RFA7367F and YFP9451R (Table S2).

### Synchronization in G1 phase with α-factor and HU-arrest

Strains SC765, Y1050, Y1051, Y1052 and Y1023 were grown until the OD600 reached 0.4. Next, cells were harvested and resuspended in fresh medium supplemented with α-factor (4 µg/mL). Additionally, α-factor (4 µg/mL) was added after 60–90 min incubation. After 2–3 h of growth at 23°C or 30°C cells were harvested and washed three times with sterile water and then resuspended in fresh SD medium and released into a new cell cycle at 23°C or 37°C (strains SC765, Y1051, Y1052 and Y1023) and at 30°C or 18°C (strains SC765 and Y1051); At indicated time points, 1 mL samples were taken. Cell pellets were resuspended in 1 mL of 70% ethanol. For HU-arrest experiments, after synchronization with α-factor, cells were released from the G1 block in the SD medium with hydroxyurea (200 mM) (SIGMA) at 23°C (strains SC765, Y1051, Y1052 and Y1023) and at 30°C (strains SC765 and Y1051) for 120 min. Next, cells were washed with sterile water and resuspended in the SD medium for incubation at 23°C or 37°C (strains SC765, Y1051, Y1052, and Y1023) and at 30°C or 18°C (strains SC765 and Y1051); At indicated time points, 1 mL samples were taken. Cell pellets were resuspended in 1 mL of 70% ethanol.

### Flow cytometry analysis

Samples were prepared for flow cytometry, as described previously [33], with modifications according to specific strain requirements. Yeast cells were stained using SYTOX Green (0.5 µM) (Invitrogen, Carlsbad, CA, USA). The DNA content was determined by measuring the SYTOX Green fluorescence signal (FL1) using Becton Dickinson FACSCalibur and CellQuest software (BD Bioscience, San Jose, CA, United States). Further analyses were done using Flowing Software (https://flowingsoftware.com/).

### Measurement of spontaneous mutation rates

To determine spontaneous mutation rates, 10 to 20 cultures of 2 or 3 independent isolates of each strain were inoculated in 2,5 mL of liquid SD medium, supplemented with the required amino acids and nitrogenous bases. Cultures were grown at 23°C or 30°C until they reached the stationary phase. Then, cells were collected by centrifugation, washed, and resuspended in 0.8% NaCl. Aliquots of concentrated cultures and their appropriate dilutions were plated on selective (containing L-canavanine) and nonselective media. Colonies were counted after 5-7 days of growth at 30°C or 23°C. Mutation rates were calculated using the MLE MUtation Rate calculator (mlemur) [34].

### Mutation spectra

A total of 192 cultures of 2 independent isolates of the Y1050 strain and 170 cultures of 2 independent isolates of the Y1060 strain were inoculated in 1 mL of liquid SD medium supplemented with the required amino acids and nitrogenous bases, lacking leucine, and grown at 30 °C. When cultures reached the stationary phase, appropriate dilutions were plated on a solid SD medium supplemented with L-canavanine. After five days of incubation at 30°C, total DNA from a single CAN^R^ colony from each culture of Y1050 and Y1060 strains was isolated and used for PCR amplification of the *CAN1* locus with primers MGCANFF and MGCANRR followed by DNA sequencing with primers Can_1666, Can_1963, Can_2241, and Can_2465 (Table S2). This enabled the identification of mutations in the *CAN1* gene from CAN^R^ cells. For statistical analysis, a contingency table and the χ2 test were used.

### Identification of Rfa1 foci by fluorescence microscopy

Cells with *RFA1–YFP* fusion were cultured at 23°C or 30°C in the SD medium supplemented with the required amino acids and nitrogenous bases, excluding leucine, until they reached the exponential growth phase before samples for fluorescent microscopy were taken. Images were taken using the Axio Imager M2 fluorescence microscope with the AxioCam MRc5 Digital Camera (Zeiss, Oberkochen, Germany) and analyzed with Axio Vision 4.8 software. The number of cells and Rfa1 foci in the cells was counted. Eight biological replicates were analyzed, with over 1700 cells for each genotype. For statistical analysis, contingency tables and χ^2^ test were applied.

## Results

### DNA replication defects in *CDC45* mutant cells

We constructed mutant derivatives of the haploid ΔI(-2)I-7B-YUNI300 *Saccharomyces cerevisiae* strain with chromosomal *cdc45-1*, *cdc45-25*, *cdc45-26,* and *cdc45-35* alleles (Table S1). Since previous studies have shown that these replication mutants are unable to grow at specific temperatures, the resulting strains were tested for growth at 18°C, 23°C, 30°C and 37°C (Figure S1). The strain carrying the *cdc45-1* allele did not grow at 18°C, consistent with previous studies [21]. Yeast cells carrying the *cdc45-25*, *cdc45-26,* and *cdc45-35* alleles could not grow at 37°C (Figure S1).

In the first step, using flow cytometry, we analyzed the DNA content profile of *cdc45* mutant strains at permissive temperatures of 30°C for *cdc45-1* and 23°C for *cdc45-25*, *cdc45-26,* and *cdc45-35* cells. This approach enables estimating the relative cell number of cells in the cell cycle’s G1, S, and G2/M phases. The results presented in Figure 2 show that in an asynchronous population of wild-type cells at 30°C, only about 10% of cells are in the S phase, while in the *cdc45-1* mutant, under the same conditions, more than 25% of cells are synthesizing DNA (Figure 2 A and B). At 23°C, about 13% of cells are in the S phase in a population of asynchronous wild-type cells. In contrast, mutant cells with *cdc45-25*, *cdc45-26,* or *cdc45-35* alleles, DNA replicating cells, constitute 25-31% of the population (Figure 2 C and D).

**Figure 2.**
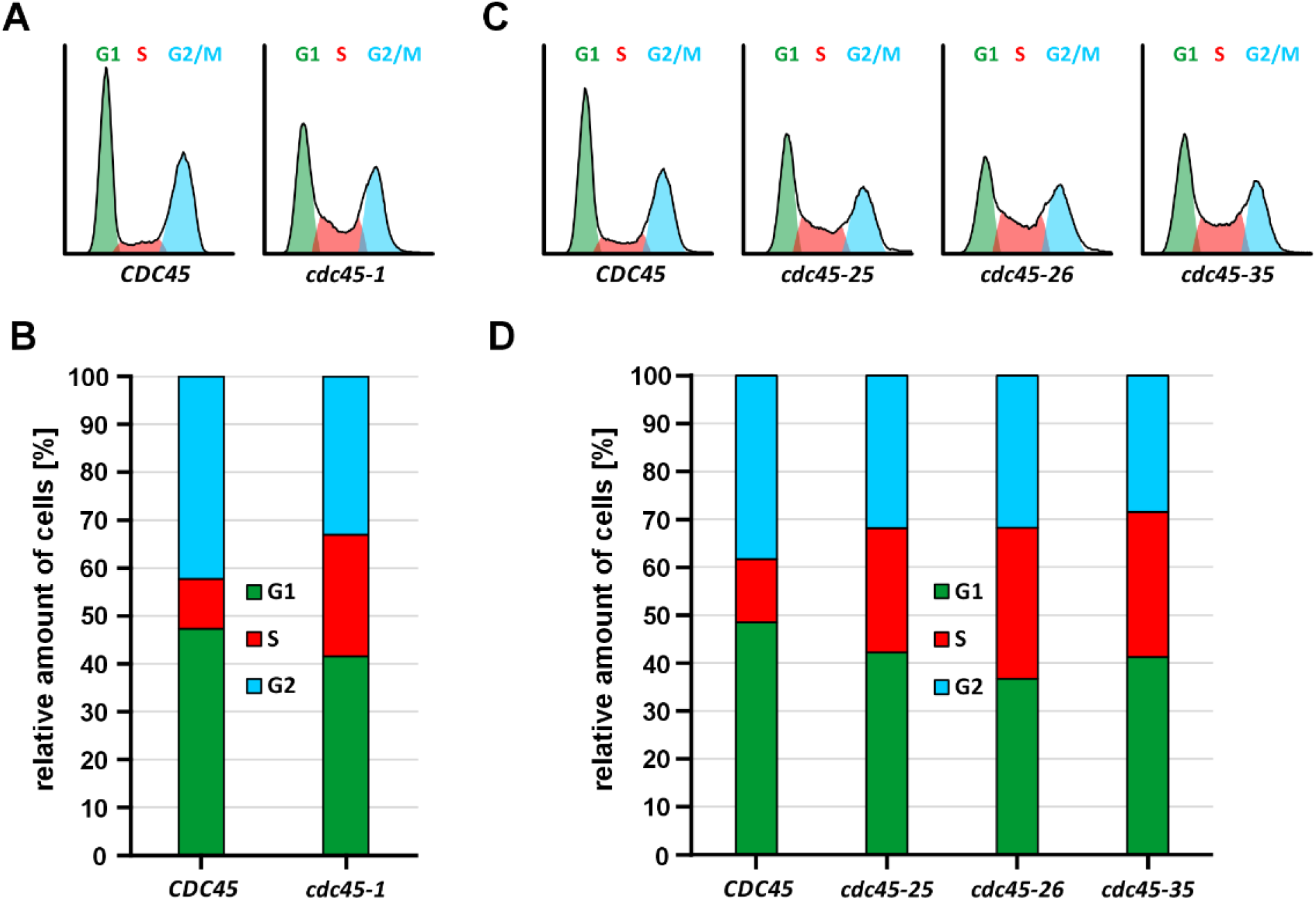
Flow cytometry analysis of DNA content in *CDC45* mutant cells in permissive conditions. DNA content was analyzed in asynchronous populations of *cdc45-1* mutant at 30°C **(A)** or *cdc45-25*, *cdc45-26*, and *cdc45-35* at 23°C **(C)**. The DNA content is shown on the x-axis, and the cell count is shown on the y-axis. The average percentage of cells in either cell cycle phase calculated for four independent repeats are shown in **(B)** and **(D)**. The statistical significance of differences between mutant and wild-type strains was assessed using a contingency table. The chi-square statistics were 735.2659, 480.9848, 904.3640, and 823.8612 for *cdc45-1*, *cdc45-25*, *cdc45-26*, and *cdc45-35*, respectively, with *p*-value <0.00001.

Next, we analyzed the cell cycle progression of cells with *cdc45-1*, *cdc45-25*, *cdc45-26*, and *cdc45-35* alleles and compared with the wild-type strain. Yeast cultures were first synchronized in the G1 phase and released into a new cell cycle at a permissive temperature (Figure 3). The entry into a new cell cycle, i.e., into the S phase, was monitored using flow cytometry-based analysis of DNA content. At 30°C, wild-type cells started DNA synthesis 45 min. after release from the G1 block and reached G2/M DNA content after 90 minutes (Figure 3A). In contrast, cells with *cdc45-1* mutation started DNA synthesis after 60-75 minutes and duplicated the genome only after 120 minutes (Figure 3A).

**Figure 3.**
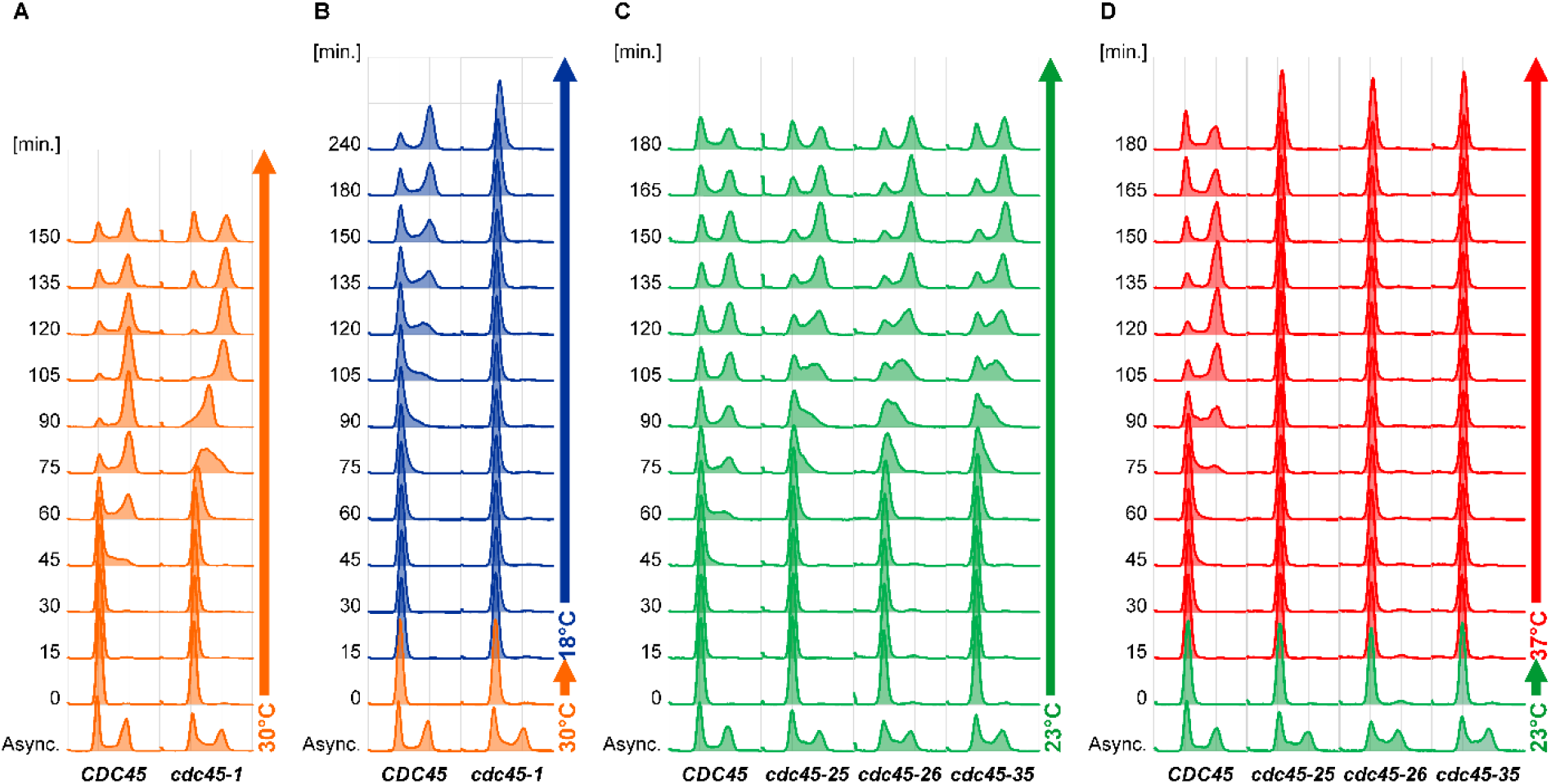
Cell cycle progression of *cdc45-1* (A, B) and *cdc45-25*, *cdc45-26*, *cdc45-35* (C, D) mutant cells. DNA content was analyzed in cells synchronized in the G1 phase and released into a new cell cycle. Cells were synchronized at the permissive temperature and released into a new cell cycle at the same temperature (30°C in **A** and 23°C in **C**) or restrictive temperature (18°C in **B** and 37°C in **D**). Samples were taken at indicated time points.

At 23°C, DNA replication in wild-type cells started 45-60 minutes after release from the G1 block and terminated after 120-135 minutes. In contrast, in *cdc45-25*, *cdc45-26*, and *cdc45-35* mutant cells, DNA synthesis started after 75 minutes (Figure 3C). After slow progression through the S phase (see time points 90, 105, and 135 minutes), mutant cells reached the G2 DNA content after 150-165 minutes (Figure 3C).

Temperature-sensitivity of replication mutants results from their inability to replicate DNA at the restrictive temperature. Therefore, we verified whether *CDC45* mutant cells can synthesize DNA at the restrictive temperature in the first cell cycle after the temperature switch. For this purpose, yeast cells with mutated *CDC45* alleles were synchronized in the G1 phase at the permissive temperature (30°C for *cdc45-1* and 23°C for *cdc45-25*, *cdc45-26*, and *cdc45-35*) and released into a new cell cycle at the restrictive temperature (18°C for *cdc45-1* and 30°C for *cdc45-25*, *cdc45-26*, and *cdc45-35*) (Figure 3). At 18°C, wild-type cells started DNA synthesis started after 75- 90 minutes and terminated after an additional 150 minutes (Figure 3B). In the same conditions, DNA content *cdc45-1* in mutant cells remained unchanged even 240 minutes after release from the G1 phase (Figure 3B). At 37°C, in wild-type cells, DNA synthesis started 75 minutes after release from the G1 phase and finished 60 minutes later. In contrast, *cdc45-25*, *cdc45-26*, and *cdc45-35* mutant cells failed to start DNA replication even after 180 minutes from G1 block release (Figure 3D).

These results show that all the studied *CDC45* mutants encounter severe problems with DNA replication. After the new cell cycle entry, *CDC45* mutant cells start DNA synthesis later than wild-type cells. DNA replication takes more time, resulting in a higher number of cells in the S-phase.

Cdc45 is recruited to the origin of DNA replication at the initiation step but is also involved in the elongation of the growing DNA strand [29]. Therefore, abnormal progression through the S phase of the cell cycle could result either from defective initiation of DNA replication or inefficient replisome activity at the elongation step. To verify this, we synchronized the cells in the G1 phase, but before releasing them into a new cell cycle, this time, they were treated with hydroxyurea (HU) for 120 minutes. This compound inhibits the ribonucleotide reductase and limits dNTPs synthesis, which stalls DNA synthesis but does not affect the initiation steps of DNA replication and replisome assembly. At 30°C, wild-type cells completed DNA synthesis 45 minutes after release from the HU block, while *cdc45-1* mutant cells did it after 60-75 minutes (Figure 4A). At 18°C, wild-type cells reached G2-specific DNA content after 105-120 minutes.

**Figure 4.**
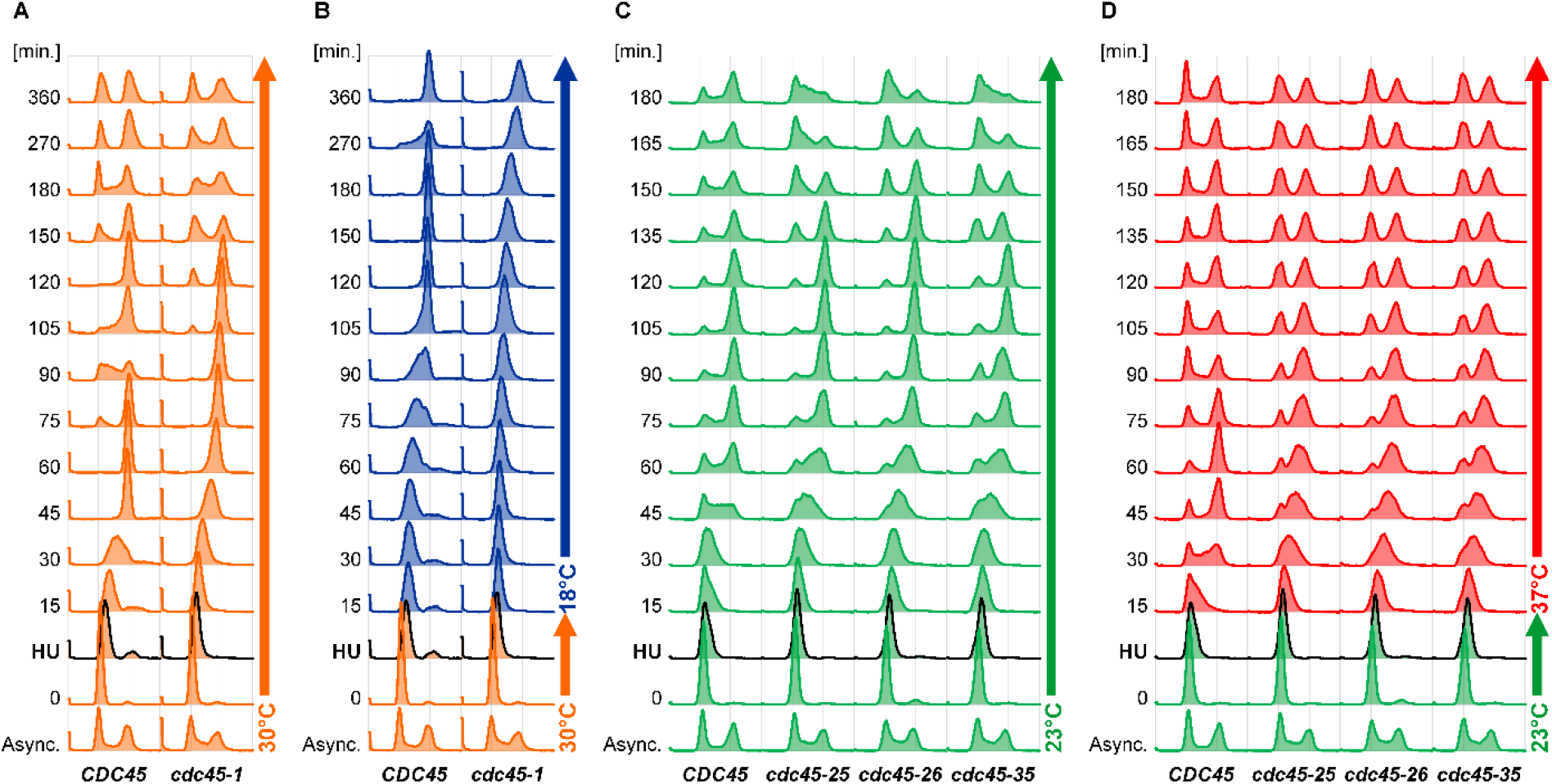
Cell cycle progression of *cdc45-1* (A, B) and *cdc45-25*, *cdc45-26*, and *cdc45-35* (C, D) mutant cells after synchronization in G1 phase and HU block (black line) at permissive temperature (30°C in A, B and 23°C in C, D) allowing initiation of DNA replication. Subsequently, cells were released into a new cell cycle at the permissive (30°C in **A** and 23°C in **C**) or restrictive (18°C in **B** and 37°C in **D**) temperature. Samples were taken at indicated time points, and DNA content was analyzed using flow cytometry. More detailed results for *cdc45-1* and the control wild-type strain, with all time points analyzed, are shown in Figure S2.

In contrast, *cdc45-1* mutant cells could not synthesize DNA efficiently and duplicated the genome after 360 minutes (Figure 4B and S2), demonstrating severe defects in DNA replication despite initiating the process at a permissive temperature. Interestingly, when the same experiment was performed with only 30 minutes of exposure to HU at the permissive temperature of 30°C, wild-type and mutant cells needed about 15-30 minutes more time to reach the G2-specific DNA content (Figure S2). At 18°C, wild-type cells reached the same DNA content over 45 minutes later than at 30°C, while *cdc45-1* mutant cells didn’t complete DNA synthesis even after 360 minutes (Figure S2).

At 23°C, wild-type and *cdc45-25*, *cdc45-26*, and *cdc45-35* mutant cells reached G2- specific DNA content 105-120 minutes after release from HU block (Figure 4C). At 37°C, wild-type cells completed DNA synthesis 60 minutes after release from HU (Figure 4D). Under the same conditions, *cdc45-25*, *cdc45-26*, and *cdc45-35* progressed slowly through the S phase and reached G2-DNA content 75-90 minutes after HU treatment (Figure 4D). This demonstrates that DNA replication initiation at the permissive temperature enables efficient DNA synthesis in *cdc45-25*, *cdc45-26*, and *cdc45-35* mutant cells, even under stress conditions.

### *CDC45* mutants accumulate single-stranded DNA

Defective replisome functioning may result in discontinued DNA synthesis at one or both DNA strands and the formation of single-stranded DNA (ssDNA) stretches. They can subsequently be used as templates for DNA synthesis by other polymerases (e.g., Pol ζ) or as substrates for homologous recombination (HR). To verify whether this is the case in the four studied *cdc45* mutants, we analyzed foci formation by Rfa1, a subunit of RPA (Replication Protein A) that binds ssDNA. For their visualization, we used a fusion protein Rfa1-Yfp [32]. Using fluorescent microscopy, we calculated the number of cells with single or multiple foci in *cdc45* mutants and control wild-type strains under permissive conditions (Figure 5). We found that in the cold-sensitive *cdc45-1* cells, over 70% of cells formed Rfa1 foci, including over 20% with double foci and over 30% with three or more foci (Figure 5). Under the same conditions (30°C), less than 63% of wild-type cells contained Rfa1 foci, with only 36% showing more than one foci. The results obtained for the three temperature-sensitive mutants *cdc45-25*, *cdc45-26*, and *cdc45-35* show 53-61% cells with Rfa1 foci, including 32-40% cells with two or more foci. In the control wild-type strain, we observed only 40% of cells with Rfa1 foci, including only 20% with two or more foci (Figure 5). Together, these results show increased ssDNA formation in all tested *CDC45* mutants.

**Figure 5.**
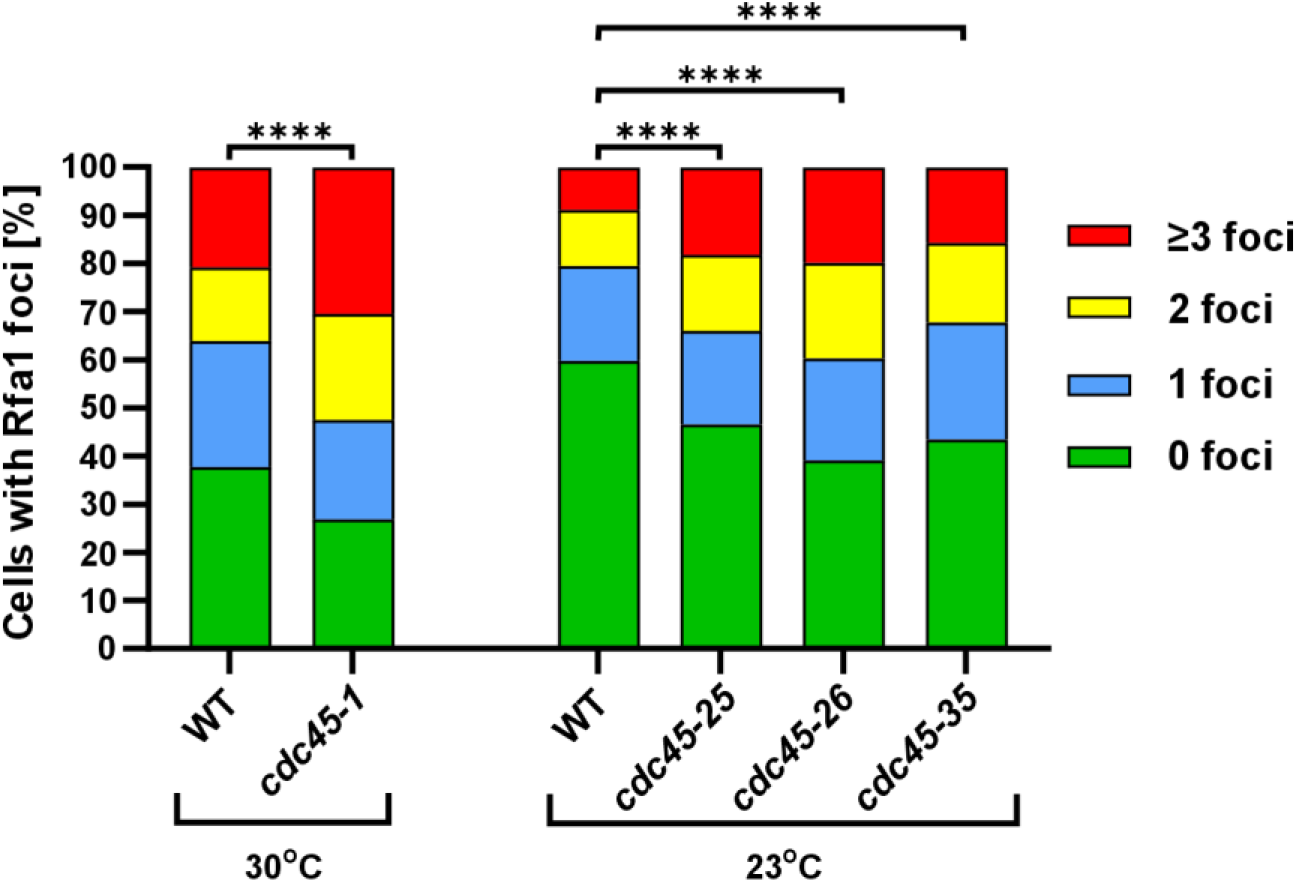
Visualization of Rfa1-Yfp foci in *cdc45-1*, *cdc45-25*, *cdc45-26*, and *cdc45-35* mutant cells. At least 200 cells were analyzed for each of the eight replicates of a given strain (over 1700 in total). A contingency table was used to calculate statistical significance.

### *CDC45* mutants demonstrate increased mutation rates partially dependent on Pol ζ

To analyze forward spontaneous mutagenesis levels in *CDC45* mutant cells, we used the *CAN1* locus [35]. For each mutant, experiments were done at the permissive temperature. At 30°C, mutation rates for wild-type cells were 101×10^-8^ and increased to over 175×10^-8^ in *cdc45-*1 mutant cells (Figure 6A). At 23°C, mutation rates in wild-type cells were 51×10^-8^ and increased to over 84×10^-8^ for *cdc45-25, cdc45-26,* and *cdc45-35* (Figure 6A). Together, these results show that the rate of spontaneous mutagenesis at the *CAN1* locus in the *cdc45* mutant strains is significantly increased compared to wild-type strains (*p*-value in Table S3), demonstrating that Cdc45 dysfunctions enhance spontaneous mutagenesis.

**Figure 6.**
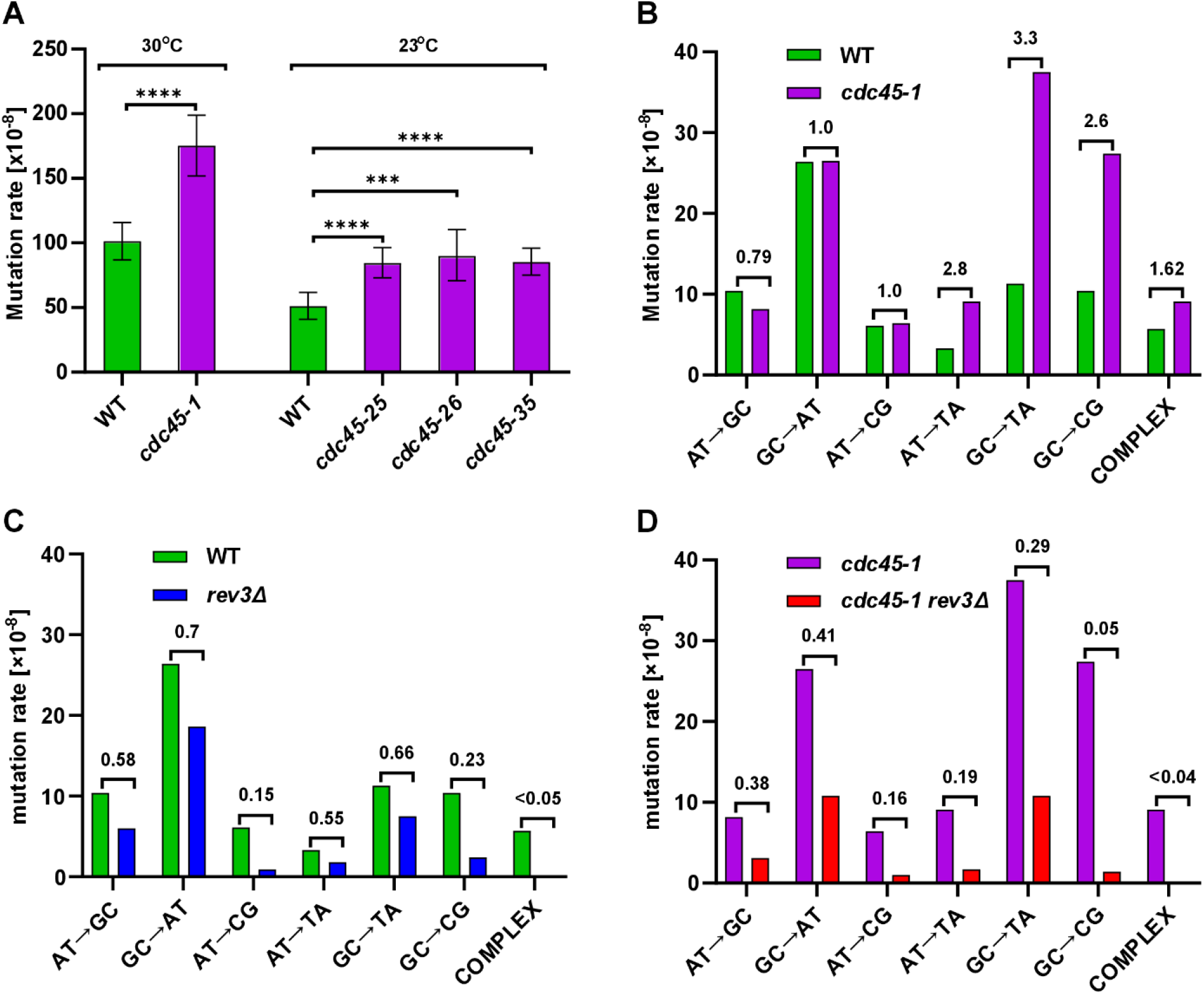
Spontaneous mutagenesis in the *cdc45* strains. **A.** Spontaneous mutation rates measured in the *cdc45* strains. The presented values were calculated using the maximum likelihood estimate mutation rate calculator (mlemur) [34]. 95% confidence intervals were calculated from at least ten independent cultures. *p*-values were adjusted using Benjamini-Hochberg correction *p*-value ≤ 0.0001 (****); ≤0.001 (***) and are shown in Table S3. **B-D.** Mutation rates with indicated fold change, calculated for specific mutation types in the *CAN1* sequence in yeast strains with the following genotypes: **B.** wild-type and *cdc45-1* strains; **C.** wild-type and *rev3*Δ strains; **D.** *cdc45-1* and *cdc45-1 rev3*Δ strains. Mutation spectra for wild-type and *rev3*Δ strains were obtained previously [59].

To obtain more details on the specificity of increased mutagenesis in *cdc45-1* mutant cells, we analyzed the sequence of the *CAN1* gene in yeast cells from canavanine-resistant colonies (Table 1 and Figure 6) . In *cdc45-1* cells, we observed a substantial increase in the frequency of three classes of transversions: GC→TA (3.3×), AT→TA (2.8×), and GC→CG (2.6×) (Figure 6B). Moreover, previous studies [36–39] and our results obtained for the wild-type strain and *rev3*Δ strain presented in this work (Figure 6C) show that the GC→CG, GC→TA, and AT→TA transversions are specific for Pol ζ.

**Table 1.** Mutation rates are calculated for specific mutation types in the *CAN1* sequence. Mutation spectra for wild-type and *rev3*Δ strains were presented previously [59].

|  | WT |  | *cdc45-1* |  | *rev3Δ* |  | *cdc45-1 rev3Δ* |  |
| --- | --- | --- | --- | --- | --- | --- | --- | --- |
| **Base substitutions** | 67.9<sup>a</sup> | 144<sup>b</sup> | 115.1 | 126 | 37.2 | 124 | 28.9 | 83 |
| **Transitions** | 36.8 | 78 | 34.7 | 38 | 24.6 | 82 | 13.9 | 40 |
| **AT→GC** | 10.4 | 22 | 8.2 | 9 | 6.0 | 20 | 3.1 | 9 |
| **GC→AT** | 26.4 | 56 | 26.5 | 29 | 18.6 | 62 | 10.8 | 31 |
| **Transversions** | 31.1 | 66 | 80.4 | 88 | 12.6 | 42 | 15.0 | 43 |
| **AT→CG** | 6.1 | 13 | 6.4 | 7 | 0.9 | 3 | 1.0 | 3 |
| **AT→TA** | 3.3 | 7 | 9.1 | 10 | 1.8 | 6 | 1.7 | 5 |
| **GC→TA** | 11.3 | 24 | 37.5 | 41 | 7.5 | 25 | 10.8 | 31 |
| **GC→CG** | 10.4 | 22 | 27.4 | 30 | 2.4 | 8 | 1.4 | 4 |
| **Indels** | 27.8 | 59 | 51.2 | 56 | 27.9 | 93 | 30.3 | 87 |
| **+1** | 5.2 | 11 | 6.4 | 7 | 3.3 | 11 | 3.5 | 10 |
| **+2** | 0.5 | 1 | 8.2 | 9 | 0.3 | 1 | 7.3 | 21 |
| **+ ≥3** | 1.9 | 4 | 5.5 | 6 | 5.1 | 17 | 5.9 | 17 |
| **− 1** | 12.3 | 26 | 24.7 | 27 | 7.8 | 26 | 8.7 | 25 |
| **− 2** | 5.2 | 11 | 5.5 | 6 | 6.3 | 21 | 3.5 | 10 |
| **− ≥3** | 2.8 | 6 | 0.9 | 1 | 5.1 | 17 | 1.4 | 4 |
| **Complex** | 5.7 | 12 | 9.1 | 10 | 0.0 | 0 | 0.0 | 0 |
| **TOTAL** | 101.3 | 215 | 175.4 | 192 | 65.0 | 217 | 59.3 | 170 |
| μ95%− | 86.9 |  | 151.8 |  | 49.1 |  | 50.8 |  |
| μ95%+ | 115.8 |  | 198.8 |  | 81.2 |  | 67.9 |  |

<sup>a</sup> Mutation rates [ $\text{Can}^R \times 10^{-8}$ ] for specific mutation types
<sup>b</sup> Number of events identified for given classes

Therefore, we also analyzed mutation spectra in *cdc45-1* cells with inactivated *REV3* gene encoding the catalytic subunit of Pol ζ. The mutation rate in the *cdc45-1 rev3*Δ strain was 65×10^-8,^ demonstrating a significant decrease compared to 175×10^-8^ in *cdc45-1 REV3* (Table 1 and Figure 7A). Rates for specific types of substitutions dropped 20× for GC→CG, over 6× for AT→CG, over 5× for AT→TA, and over 3× for GC→TA (Figure 6D). Additionally, the mutation rate for complex mutations, which are also characterized as characteristic of Pol ζ dropped over 26× in in *cdc45-1 rev3*Δ cells when compared to *cdc45-1* (Table 1).

**Figure 7.**
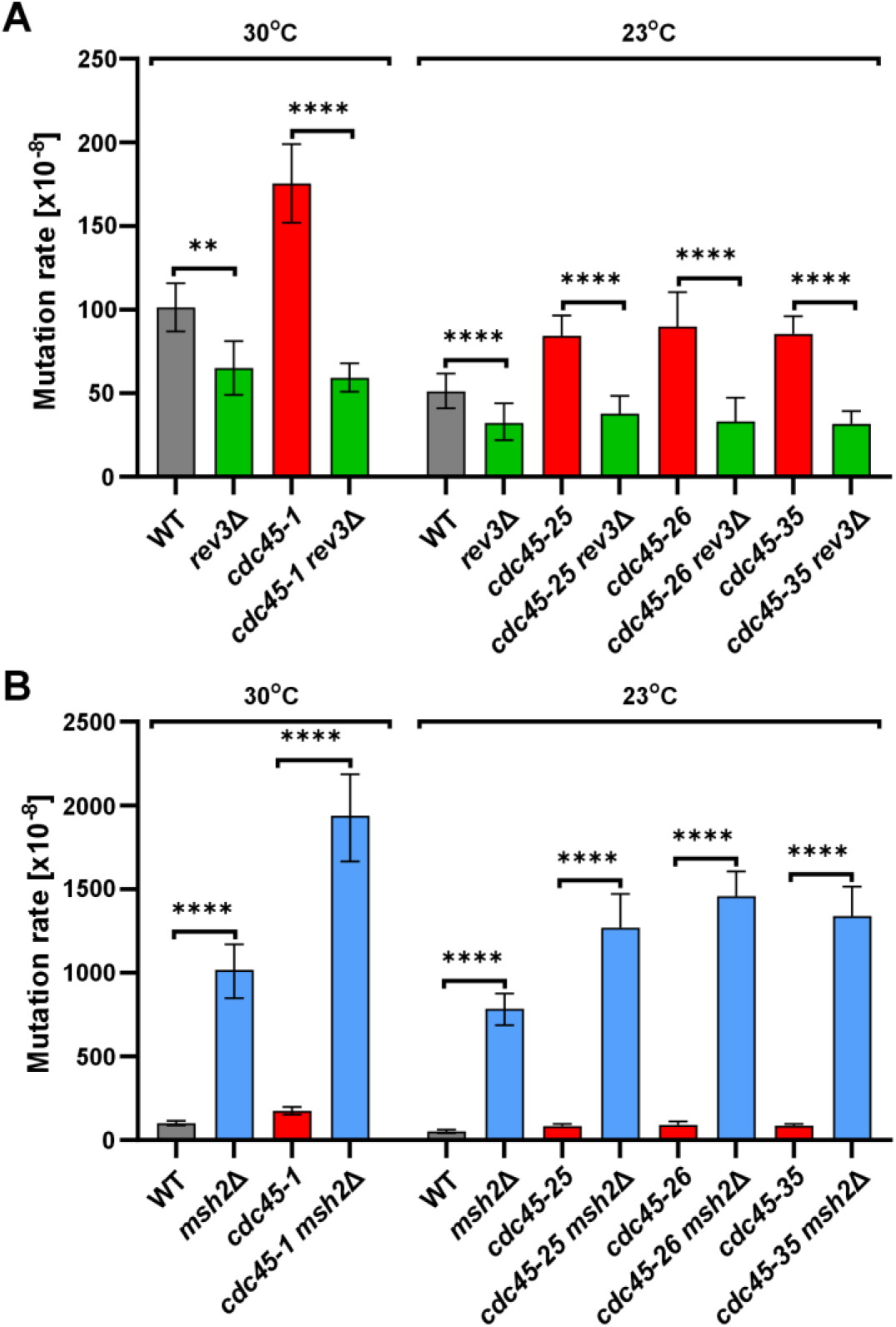
Spontaneous mutation rates were measured in *cdc45* mutant strains with *rev3*Δ (A) and *msh2*Δ (B). The presented values were calculated using the maximum likelihood estimate mutation rate calculator (mlemur) [34]. 95% confidence intervals were calculated from at least ten independent cultures. The *p*-values were adjusted using the Benjamini-Hochberg correction. *p*-value ≤ 0.0001 (****); ≤0.01 (**). The *p*-values are shown in Table S3.

The involvement of Pol ζ in increased mutagenesis was also tested in *cdc45-25*, *cdc45-26*, and *cdc45-35* cells using strain derivatives with deletion of *REV3* (Figure 7A). In all three cases, we observed that in comparison to respective *REV3* controls, mutation rates significantly dropped in *cdc45-25 rev3*Δ, *cdc45-26 rev3*Δ, and *cdc45-35 rev3*Δ (by 55%, 63%, and 62%, respectively) (Figure 7A). This result is similar to the one observed for the *cdc45-1 rev3*Δ strain (66%) (Figure 7A). These results demonstrate that Pol ζ significantly contributes to increased mutation rates in *cdc45* mutant cells.

To analyze whether *cdc45* mutations decrease replication fidelity, we inactivated the mismatch repair (MMR) mechanism involved in the repair of base-base mismatches and small insertion-deletion loops made by replicative polymerases [40–42]. This system, temporally coupled with DNA replication, does not correct errors made during translesion DNA synthesis (Pol ζ), DNA repair, or recombination [43]. To inactivate MMR, we deleted the *MSH2* gene encoding the Msh2 protein, which, together with Msh6, binds to the mismatch to initiate the repair mechanism. At 30°C, when compared to the wild-type strain, mutation rates in *msh2*Δ cells increased 10-fold from 101 to 1018×10-8 (Figure 7B). In *cdc45-1* cells, the inactivation of *MSH2* increased mutation rates 11-fold from 175 to 1936×10-8 (Figure 7B). At 23°C, mutation rates in *msh2*Δ cells increased 15-fold compared to wild-type cells (from 51 up to 784×10^-8^, Figure 7B). Similarly, in *cdc45-25*, *cdc45-26*, and *cdc45-35* cells, deletion of *MSH2* increased mutation rates over 15-fold from 84, 89, 85×10^-8^ up to 1269, 1457, and 1338×10^-8^, respectively (Figure 7B). Together, these results demonstrate that the vast majority of errors occurring in studied *cdc45* mutants are replication errors corrected by MMR

## Discussion

Chromosomal DNA replication requires the coordinated action of the CMG helicase and major DNA polymerases. The correct progression of DNA replication and high fidelity of this process is ensured not only by the major DNA polymerases but also by the non-catalytic elements of the replisome [33,38,39,44–47]. The Cdc45 is an essential protein, which, together with the GINS complex and the Mcm2-7 heterohexamer, forms the active CMG helicase which plays a key role during chromosomal DNA replication. Studies of the non-catalytic Cdc45 protein are of growing interest because its defect leads to the development of Meier-Gorlin syndrome (MGS) [22–24], craniosynostosis [48], or cancer [49,50]. For example, Meier-Gorlin syndrome is an autosomal recessive disease characterized by severe intrauterine and postnatal growth retardation, microcephaly, bilateral microtia, aplasia or hypoplasia of the patellae [51,52]. In general, mutations leading to MGS are biallelic and were identified in pre-RC-encoding genes [53]. In Cdc45, these mutations result in amino acid substitutions or alter splicing [23]. Variants of the remaining *CDC45* allele in patients with 22q11.2 deletion were also identified as a causative factor of various anomalies, including craniosynostosis, though independent of the Meier-Gorlin syndrome [48]. The effects of Cdc45 dysfunctions were thought to affect the rate of DNA replication and thus, cell proliferation. However, as it has been shown in the yeast model, Cdc45 is also a targeting factor for Rad53, the main replication checkpoint kinase. After replication checkpoint activation, phosphorylation of Cdc45 by Rad53 results in their binding and subsequent phosphorylation of Sld3 by the same kinase. This, in turn, inhibits the interaction of Cdc45 with Sld3, enabling its reactivation after checkpoint inactivation [22]. Besides binding to helicase subunits, Cdc45 also interacts with the catalytic domain of Pol2, the main subunit of Pol ε [54,55].

The previous studies analyzing *CDC45* mutants focused on the cold-sensitive *cdc45-1* mutant [14,56,57]. First, it has been demonstrated that when the incubation temperature for *cdc45-1* is switched from 30°C (permissive) to 15°C (restrictive), mutant cells accumulate partially replicated DNA [14]. When the same temperature shift was done after cell synchronization in the G1 phase, *cdc45-1* could not enter the S phase for 2-3 hours [14]. In another study published in the same year, the same assay was done with a temperature shift to 11°C, again resulting in the accumulation of cells with 1C DNA content [14]. In the present work, we used 18°C as the restrictive temperature and observed a similar effect (Figure 3B). Interestingly, we found that after the G1 block, *cdc45-1* mutant cells failed to start DNA replication even four hours after release into a new cell cycle at 18°C. In contrast, wild-type cells completed DNA replication under the same conditions. Three new *CDC45* mutants presented in this work, i. e., *cdc45-25*, *cdc45-26*, and *cdc45-35,* are temperature-sensitive, i.e., their growth is inhibited at 37°C. Thus, they were grown and synchronized with α-factor at 23°C (permissive temperature) and released from the G1 block at 37°C (restrictive temperature). All three mutants were unable to start DNA replication at 37°C, while the wild-type strain, under the same conditions, completed DNA replication after 120 minutes (Figure 3D). Together, these results demonstrate important defects of DNA replication in cells with *CDC45* mutations. An alternative for conditional *CDC45* mutants came with the heat-inducible degron mutant *cdc45-td*, which enables the inactivation of the protein at a specific time [58]. The advantage of this approach is that the protein is fully functional at permissive temperatures. However, this cannot reflect the physiological conditions of cells with amino acid substitutions in protein regions with specific functions (e.g., interaction with other proteins). Nevertheless, the complete inactivation of *CDC45* after cell synchronization in the G1 phase resulted in cell stacking in the G1/early S phase with no DNA synthesis [58]. To verify whether Cdc45 defects affect DNA replication initiation or elongation step, one can use HU to block DNA synthesis without affecting replication initiation. When *cdc45-td* cells, after synchronization with α-factor, were transiently incubated with HU at a permissive temperature, heat-induced degradation of Cdc45 resulted in incomplete DNA replication [58]. This demonstrated that Cdc45 is also involved in the elongation step of DNA replication. In the present work, we observed that the three studied temperature-sensitive mutants (*cdc45-25*, *cdc45-26*, and *cdc45-35*), after DNA replication initiation at specific permissive temperatures, followed by release at restrictive temperature, performed DNA replication almost as fast as in wild-type cells (Figure 4D). However, *cdc45-1* mutant cells showed prolonged entry into the S phase, requiring nearly three times more time than wild-type cells to complete DNA replication (Figure 4B). Importantly, when the time for DNA replication initiation was reduced to 30 min., *cdc45-1* cells needed even more time to complete DNA synthesis, and their DNA content didn’t reach G2 even after six hours. Our result for *cdc45-1* is similar to the inability to complete DNA replication observed as an effect of induced Cdc45 degradation [58]. Together, we interpret the results obtained in this work as showing that *cdc45-25*, *cdc45-26*, and *cdc45-35* mutants are defective in replication initiation since HU-block enables rapid completion of DNA replication. In contrast, *cdc45-1,* despite initiation-favorable conditions, fails to properly proceed with the DNA replication elongation step (Figure 4B), which does not support the conclusions drawn from the data discussed, yet not shown in a previous study on *cdc45-1* [14].

Our results also demonstrate that DNA synthesis is defective in *CDC45* mutant cells, even at permissive temperatures. Clearly, both cold- and temperature-sensitive mutants require slightly more time to complete DNA synthesis after S-phase entry (Figure 3A and C). This conclusion is consistent with another result obtained in this work, showing an increased frequency of ssDNA formation (Figure 5), a consequence of defective replisome functioning. These defects result in increased mutation rates, which almost doubled in *cdc45-1* mutant cells and increased by about 1.6× in *cdc45-25*, *cdc45-26*, and *cdc45-35* mutant cells (Figure 6A and Table S3). To our knowledge, this is the first report showing increased mutation frequencies in *CDC45* mutant cells. In mismatch repair proficient strains, this increase is mainly due to the activity of Pol ζ. As shown in Figure 7A, the deletion of *REV3* significantly reduces the mutation rate in the four *CDC45* mutant cells to the level observed in respective control *rev*Δ strains (Table S3). In parallel, our analysis of mutation spectra in *cdc45-1* cells clearly shows an increased incidence of mutations previously associated with Pol ζ’s activity, i.e., GC→CG, GC→TA, and AT→TA [36–39] (Figure 6B, C and Table 1). Moreover, in *cdc45-1 rev*Δ cells, mutation rates for these substitutions dropped to the level observed in *rev*Δ cells (Figure 6C, D). Together, these results show an important role of Pol ζ in the increased mutation rates in this *CDC45* mutant in mismatch repair proficient strains.

The increased participation in the replication of undamaged DNA of the low fidelity Polζ, which elevates the rate of spontaneous mutation, may be due to a phenomenon called defective-replisome-induced mutagenesis (DRIM) [36]. Previously, we showed that DRIM can also be promoted by defects in non-catalytic replisome components [39,59,60]. The presence of ssDNA in *cdc45* mutants may suggest that under defective CMGE conditions, Pol ζ is recruited to the stalled (impaired) replication forks.

Interestingly, under conditions that allow us to visualize replication errors, e.g., under conditions of defective mismatch repair (MMR) mechanism, when the *MSH2* gene is deleted, *CDC45* mutants demonstrate a strong mutator phenotype. Compared with the strains carrying single mutations, a robust synergistic effect is observed for *CAN1* mutations in strains that combine each *CDC45* mutation with *msh2*Δs (Figure 7B and Table S3). This result demonstrates that the DNA replication mechanism introduces significantly more errors in any of the four studied mutants in *CDC45*.

Amino acid substitutions caused by the four mutations analyzed in this work may cause different defects in Cdc45 functioning. The *cdc45-1* allele, which confers cold sensitivity, encodes a G367D substitution in the CID domain interacting with the Mcm2 subunit of the DNA helicase core (Figure 1), as suggested previously [18]. The *cdc45- 1* was initially isolated as a cold-sensitive cell-division-cycle mutant [21]. Later, it has been shown that *cdc45-1* is synthetically lethal with *mcm2-1*, *mcm3-1*, and *orc2-1* mutants [14]. Among alleles conferring temperature sensitivity, *cdc45-26* encodes a W481R substitution in the Rec J fold interacting with the Psf1 subunit of GINS (Figure 1), and c*dc45-25* encodes an L131P substitution in the DHH, which was suggested to weaken the affinity of Cdc45 to Sld3 [18]. Finally, the *cdc45-35* allele encodes an S242P substitution in the helical insertion specific for Cdc45 and is absent in the bacterial RecJ protein, involved in replisome interactions, including CMG [18] and Pol2 [54]. Recently, structural studies have demonstrated that this site interacts with Sld3 [61]. The importance of Cdc45-Sld3 binding was pointed out previously as essential for their association with the origin of replication [29] and, as mentioned above, for Sld3 phosphorylation upon checkpoint activation [22]. We can postulate that interactions of Cdc45 with other proteins (Mcm, Psf1, Sld3) are essential mainly for replisome assembly, which is supported by the observation that DNA replication progression in *cdc45-25, -26, and -35* cells is almost unaffected at the restrictive temperature if only the initiation of the process occurs at the permissive temperature (Figure 4D). Therefore, the *cdc45-1* allele differs from the three others since it apparently strongly affects later steps of DNA replication at the restrictive temperature after initiating the process in permissive conditions (Figure 4B). Although the low temperature slows down DNA replication even in wild-type cells by about 60 minutes, the effect observed in *cdc45-1* cells is significantly more pronounced (Figure 4B).

Although in permissive conditions, DNA replication is only slightly delayed in all four mutants analyzed in this work, the effect of these alleles on genome stability is significant: they strongly increase the rates of replication errors (Figure 7B) but also result in incomplete DNA synthesis generating ssDNA regions. In addition, DNA replication is rescued by Pol ζ, contributing to increased mutation rates (Figures 6B-D and 7A). These results strongly support the postulated involvement of Cdc45 in DNA replication and the maintenance of genome stability.

## Acknowledgements

We are grateful to Hiroyuki Araki (NIG, Mishima, Japan) for providing *CDC45* alleles and many valuable discussions.

## Author contributions

Conceptualization (M.D-K, I. J. F., M. D.), Data curation (M.D-K, D. C.; A. K., M. J., M. D.), Formal analysis (I. J. F., M. D.), Funding acquisition (M.D-K), Investigation (M.D-K, D. C.; A. K., M. J.), Methodology (M.D-K, I. J. F., M. D.), Supervision (I. J. F., M. D.), Validation (I. J. F., M. D.), Visualization (M. D.), Writing – original draft (M. D-K., M. D.), Writing – review and editing (M. D-K., D. C.; A. K., I. J. F, M. D.)

## Supplementary materials

**Figure S1. Temperature-sensitivity of *CDC45* mutants.** Dilutions of yeast cultures with specified *CDC45* alleles were plated and incubated at indicated temperatures.

**Figure S2. Cell cycle progression of *cdc45-1* mutant cells after synchronization in G1 phase and HU block at permissive temperature (30°C) for 120 or 30 min.** Subsequently, cells were released into a new cell cycle at the permissive (30°C) or restrictive (18°C) temperature. Samples were taken at indicated time points, and DNA content was analyzed using flow cytometry.

**Table S1.** Yeast strains used in this study.

**Table S2.** Primers used in this study.

**Table S3.** *p*-values adjusted using Benjamini-Hochberg correction obtained for data presented in figure 6.

