## Supplementary materials for "The effects of *CDC45* mutations on DNA replication and genome stability"

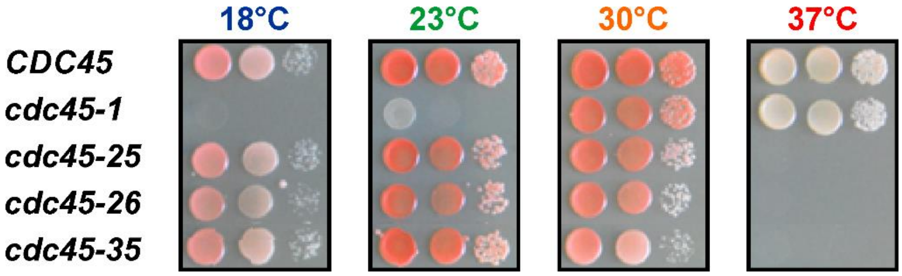


**Figure S1. Temperature-sensitivity of *CDC45* mutants.** Dilutions of yeast cultures with specified *CDC45* alleles were plated and incubated at indicated temperatures.

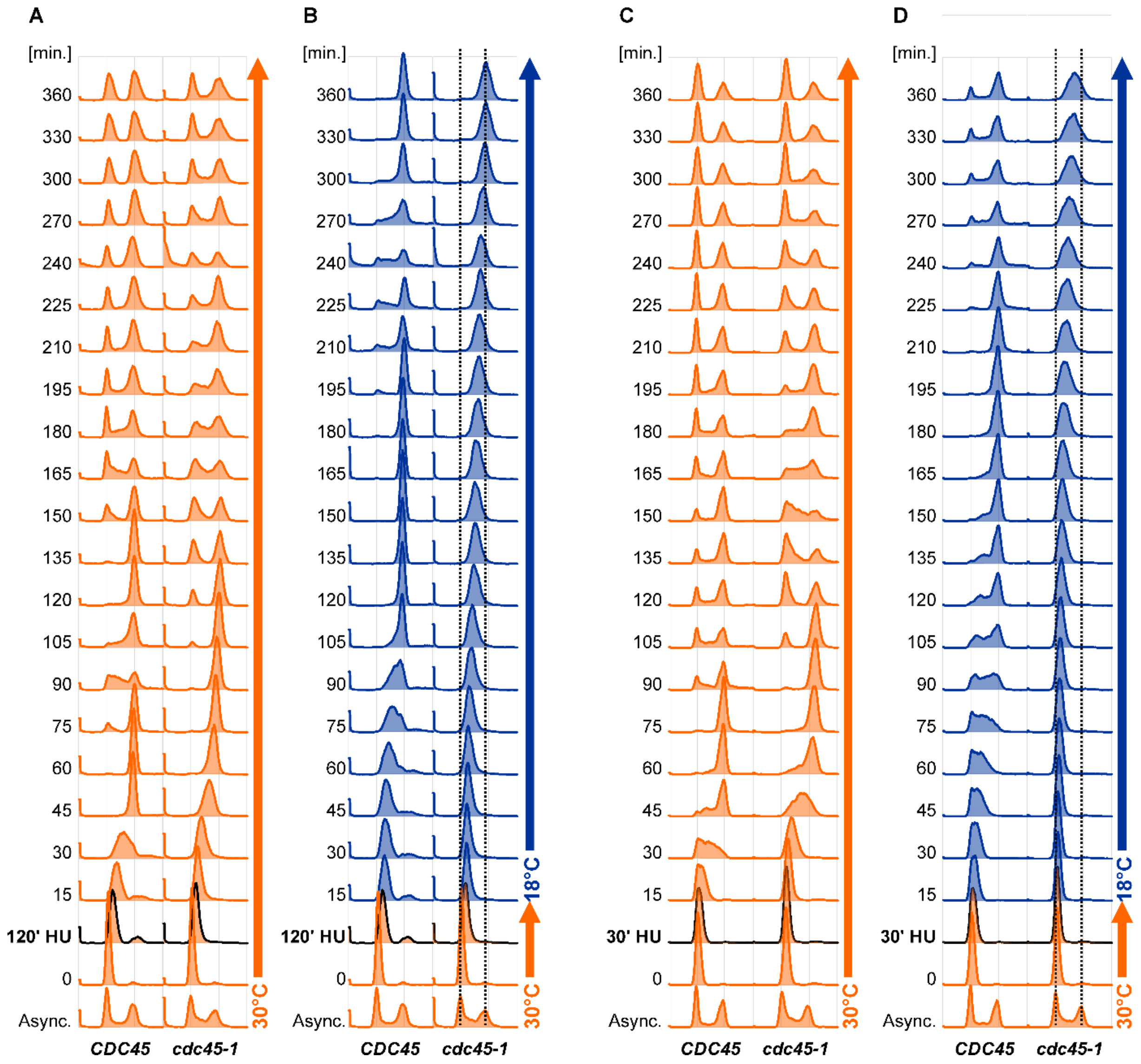


**Figure S2**. **Cell cycle progression of *cdc45-1* mutant cells after synchronization in G1 phase and HU block at permissive temperature (30°C) for 120 or 30 min.** Subsequently, cells were released into a new cell cycle at the permissive (30°C) or restrictive (18°C) temperature. Samples were taken at indicated time points, and DNA content was analyzed using flow cytometry.

**Table S1.** Yeast strains used in this study.

| Strain | Genotype | Source |
|---|---|---|
| SC765[a] | *MATa CAN1 his7-2 leu2Δ::hisG ura3Δ trp1-289 ade2-1 lys2ΔGG2899-2900* | [38] |
| Y1050 | *As SC765, but* ***cdc45-1*** | This work |
| Y1051 | *As SC765, but* ***cdc45-25*** | This work |
| Y1052 | *As SC765, but* ***cdc45-26*** | This work |
| Y1053 | *As SC765, but* ***cdc45-35*** | This work |
| Y1054 | *As SC765, but* ***msh2::NAT1*** | This work |
| Y1055 | *As SC765, but* ***msh2::NAT1 cdc45-1*** | This work |
| Y1056 | *As SC765, but* ***msh2::NAT1 cdc45-25*** | This work |
| Y1057 | *As SC765, but* ***msh2::NAT1 cdc45-26*** | This work |
| Y1058 | *As SC765, but* ***msh2::NAT1 cdc45-35*** | This work |
| Y1059 | *As SC765, but* ***rev3::LEU2*** | This work |
| Y1060 | *As SC765, but* ***rev3::LEU2 cdc45-1*** | This work |
| Y1061 | *As SC765, but* ***rev3::LEU2 cdc45-25*** | This work |
| Y1062 | *As SC765, but* ***rev3::LEU2 cdc45-26*** | This work |
| Y1063 | *As SC765, but* ***rev3::LEU2 cdc45-35*** | This work |
| Y1064 | *As SC765, but* ***(RFA1-YFP, LEU2) cdc45-1*** | This work |
| Y1065 | *As SC765, but* ***(RFA1-YFP, LEU2) cdc45-25*** | This work |
| Y1066 | *As SC765, but* ***(RFA1-YFP, LEU2) cdc45-26*** | This work |
| Y1067 | *As SC765, but* ***(RFA1-YFP, LEU2) cdc45-35*** | This work |

[a] This strain is a derivative of ΔI(-2)I-7B-YUNI300 [28]

**Table S2.** Primers used in this study.

| Primer name | Sequence 5'→3'/application |
|---|---|
| | **PCR amplification and DNA sequencing of *CAN1* locus** |
| MGCANFF | AAGAGTGGTTGCGAACAGAG |
| MGCANRR | GGAGCAAGATTGTTGTGGTG |
| Can_1666 | ATATTTGACAGGGAACAAGT |
| Can_1963 | GATGGCTCTTGGAACGGA |
| Can_2241 | TGTCAAGGACCACCAAAG |
| Can_2465 | GTAACTCGTCACGAGAGA |
| | **Gene disruptions** |
| MSH2_UPTEF | CTTTATCTGCTGACCTAACATCAAAATCCTCAGATTAAAAGTATGAGATCTGTTTAGCTTGCC |
| MSH2_DNTEF | ATTATCTATCGATTCTCACTTAAGATGTCGTTGTAATATTAATTATTCGAGCTCGTTTTCGACAC |
| | **Verification of gene disruptions** |
| MSH2 A | CGTATAAACAAAGCCAAAGACAAGT |
| MSH2 B | CCCAATTGAATCAAGAAACTCTCTA |
| MSH2 C | TGAATTGACAGAATTGTCTGAAAAA |
| MSH2 D | ACATCTCTTGTTTATCCCATCCATA |
| Msh2_up | TCGGTTCTTACTGCCAAGTG |
| Msh2_prdw | CATACAGGAGGTGATCCGGT |
| REV3 A | AATTCTGCCAATCTATTTGATCTTG |
| REV3 B | TCTGATTTAGAGGATGATCTAACCG |
| REV3 C | TAAATGAAGACCATAGAGCAGAACC |
| REV3 D | CACCAGATAGAGTTTTGAACGAAAT |
| NAT1 UO | ACCGGTAAGCCGTGTCGTCAAG |
| NAT1 DO | GCTTCGTGGTCGTCTCGTACTC |
| | **PCR amplification and DNA sequencing of *cdc45-1, cdc45-25, cdc45-26* and *cdc45-35* locus** |
| cdc45_up | AAGCCATGCGAATCCTAC |
| cdc45_dw | GCCGCGCACAAAATATGG |
| cdc45_1 | CACTAGAGAGAAGGCACATA |
| cdc45_2 | GGTACGGTGGATGACACATT |
| cdc45_3 | TCCAACCGGATTACTACCTT |
| cdc45_4 | CGTGGCATTCAACTAGCACA |
| | **Confirmation of the *RFA1-YFP* fusion cassette** |
| RFA6231R | ACGGTTCACAATCCCTACAG |
| RFA7367F | GCCGCAACGCAAACTTCATC |
| YFP9451R | CTTCGGGCATGGCACTCTTG |

**Table S3.** *p*-values adjusted using Benjamini-Hochberg correction obtained for data presented in figure 6.

| | Strains | | | ***p*-value corrected**[a] |
|---|---|---|---|---|
| 30°C | WT | vs | *cdc45-1* | **2,58E-07** |
| | WT | vs | *msh2Δ* | **2,16E-18** |
| | *cdc45-1* | vs | *cdc45-1 msh2Δ* | **8,78E-26** |
| | *msh2Δ* | vs | *cdc45-1 msh2Δ* | **1,16E-07** |
| | WT | vs | *rev3Δ* | **1,22E-03** |
| | *cdc45-1* | vs | *cdc45-1 rev3Δ* | **2,08E-18** |
| | *rev3Δ* | vs | *cdc45-1 rev3Δ* | 5,34E-01 |
| 23°C | WT | vs | *cdc45-25* | **4,82E-05** |
| | WT | vs | *cdc45-26* | **6,04E-04** |
| | WT | vs | *cdc45-35* | **1,16E-05** |
| | WT | vs | *msh2Δ* | **3,29E-38** |
| | *cdc45-25* | vs | *cdc45-25 msh2Δ* | **8,98E-21** |
| | *cdc45-26* | vs | *cdc45-26 msh2Δ* | **4,68E-48** |
| | *cdc45-35* | vs | *cdc45-35 msh2Δ* | **1,26E-30** |
| | *msh2Δ* | vs | *cdc45-25 msh2Δ* | **1,79E-04** |
| | *msh2Δ* | vs | *cdc45-26 msh2Δ* | **1,86E-11** |
| | *msh2Δ* | vs | *cdc45-35 msh2Δ* | **1,88E-06** |
| | WT | vs | *rev3Δ* | **2,07E-02** |
| | *cdc45-25* | vs | *cdc45-25 rev3Δ* | **1,43E-08** |
| | *cdc45-26* | vs | *cdc45-26 rev3Δ* | **5,75E-06** |
| | *cdc45-35* | vs | *cdc45-35 rev3Δ* | **3,62E-16** |
| | *rev3Δ* | vs | *cdc45-25 rev3Δ* | 5,31E-01 |
| | *rev3Δ* | vs | *cdc45-26 rev3Δ* | 9,28E-01 |
| | *rev3Δ* | vs | *cdc45-35 rev3Δ* | 9,28E-01 |

[a] *p*-values were calculated using Benjamini-Hochberg correction for multiple comparisons.
